# No clear monogenic links between left-handedness and *situs inversus*

**DOI:** 10.1101/422964

**Authors:** Merel C. Postema, Amaia Carrion-Castillo, Simon E. Fisher, Guy Vingerhoets, Clyde Francks

**Affiliations:** Department of Language & Genetics, Max Planck Institute for Psycholinguistics, Nijmegen, Netherlands; Donders Institute for Brain, Cognition and Behaviour, Radboud University, Nijmegen, Netherlands; Department of Experimental Psychology, Ghent University, Belgium

**Keywords:** *Situs inversus*, primary ciliary dyskinesia, left-handedness, whole genome sequencing

## Abstract

Left-handedness is a complex trait which might sometimes involve rare, monogenic contributions. *Situs inversus* (SI) of the visceral organs can occur with Primary Ciliary Dyskinesia (PCD), due to mutations which affect left-right axis formation. Roughly 10% of people with SI and PCD are left-handed, similar to the general population. However, in non-PCD SI, the rate of left-handedness may be elevated. We sequenced the genomes of nine non-PCD SI people who show an elevated rate of left-handedness (five out of nine). We also sequenced six SI people with PCD as positive controls, and fifteen unaffected people as technical controls. Recessive mutations in known PCD genes were found in all positive controls with PCD. Of the nine non-PCD SI cases, two had recessive mutations in known PCD genes, suggesting reduced penetrance for PCD, and one had a recessive mutation in the non-PCD laterality gene *PKD1L1*. However, six of the nine non-PCD SI cases, including most of the left-handers, had no mutations in likely candidate genes, nor any significant biological pathways affected by their mutations. Therefore we did not identify a molecular link between visceral and brain laterality. While we cannot exclude a monogenic basis for non-PCD SI with left-handedness, multifactorial and non-genetic models must also be considered.

## Introduction

A fundamental question in human neurobiology is how the brain becomes functionally left-right asymmetrical. For example, approximately 90% of people are right-handed and have left-hemisphere dominance for language, among other lateralized functions, but the developmental basis for this asymmetry remains unknown^1^. One possibility is that early embryonic mechanisms which give rise to asymmetries of the visceral organs also impact on brain asymmetries. However, this has not been previously addressed by genetic mutation screening in people who are both left-handed and have altered forms of visceral laterality.

Roughly 1:6,000-8,000 people have *situs inversus* (SI), a mirror reversal of the normal asymmetrical arrangement of the viscera ^2,3^. SI can occur alone or in combination with Primary Ciliary Dyskinesia (PCD), a recessive genetic disorder which involves mutations that disrupt motile cilia ^2^. Cilia are hair-like organelles that protrude from the cell surface into the extracellular space ^4^. They are expressed in various tissues including the respiratory epithelium ^5^, so that a disruption of ciliary motility can cause symptoms such as chronic bronchitis, inflamed or infected sinuses ^6^, which are often present in PCD.

Motile cilia are also expressed early in development, within an embryonic structure called the ‘node’, where they generate a leftward fluid flow that helps to create the left-right body axis ^5^. The leftward direction of the nodal flow may be explained by a posterior tilt of the cilia together with their clockwise rotation, arising ultimately from molecular chirality of their component proteins ^7-9^. When leftward nodal flow is absent due to recessive PCD-causing mutations, many of which affect component proteins of the ciliary cytoskeleton, it becomes a matter of chance whether the viscera will take up the normal or mirror-reversed positioning ^10^.

However, roughly three quarters of people with SI do not have PCD ^2^, and the causes of their SI remain largely unknown. A few genes have been reported to be involved in SI without PCD, and some of these can also cause partial disruptions of visceral laterality, known as heterotaxy or *situs ambiguus* (SA), including *ZIC3* [MIM:300265] ^11^, *CCDC11* [MIM:614759] ^12^, *WDR16* [MIM:609804] ^13^, *NME7* [MIM:613465] ^14^, and *PKD1L1* [MIM:609721] ^15^. The mechanisms by which these non-PCD, SI-causing genes influence visceral laterality are not well understood, but most code for cilia-associated proteins, rather than coding directly for cytoskeletal components of cilia.

Intriguingly, the general population rate of 85-90% right-handedness is not altered in SI people with PCD ^16,17^. This implies a developmental dissociation between brain laterality for hand motor control and nodal-ciliary visceral patterning. However, in the only study of handedness in SI to include non-PCD cases, Vingerhoets *et al*. reported that five of nine SI cases without PCD were left-handed ^18^. Although based on a small sample, this suggests developmental mechanisms which might indeed link handedness and visceral laterality, but independently of genes involved in PCD.

One study reported a possible genetic link between a continuous measure of left-versus-right hand motor skill and genes involved in visceral laterality ^19^, based on analyzing genetic variants which are common in the population. However, the sample size of under 3000 subjects was low for complex-trait genome-wide association analysis using common genetic variants. A much larger study of over 300,000 subjects from the UK Biobank found no support for a link of left-handedness to genes involved in visceral asymmetry ^20^. The same large study identified an association of a common variant at the *MAP2* gene with left-handedness, with a very small effect ^20^. Left-handedness has a heritability of roughly 25% based on twin and family data ^21^, but only around 2% based on genome-wide genotype data for common polymorphisms, within the UK Biobank dataset ^22^.

It has been proposed that left-handedness may sometimes occur due to genetic mutations which are rare in the population, but might have substantial effects on brain laterality when present ^23,24^. Rare genetic effects are not well captured in genome-wide association studies, which are focused on relatively common variation ^20^. Here we performed an exploratory whole genome sequencing study in the same set of 15 SI subjects studied by Vingerhoets *et al*., as well as 15 healthy controls matched for age, sex, education, and handedness (**Table 1**). The goal was to identify rare, highly penetrant mutations in the nine people with non-PCD SI, as they show an elevated rate of left-handedness. This approach had the potential to yield novel insights into the developmental origins of left-right patterning of both brain and body.

**Table 1.** The most likely causal recessive mutations in the fifteen SI subjects

| Subj | SI group | Sex/<br>Age | EHI | NH | CHD | Daily<br>wet<br>cough | Type | Gene | Clinvar | rs ID | Start<br>position | ref | alt | MAF | AAC | impact |
| --- | --- | --- | --- | --- | --- | --- | --- | --- | --- | --- | --- | --- | --- | --- | --- | --- |
| SI02 | non-PCD | M/50 | 0.9 | R | 0 | ? | hom | *PKD1L1* | SA \| SI | rs528302390 | 47870810 | T | TCA | 0.00109 | - | splice donor |
| SI03 | non-PCD | F/26 | -0.8 | L | 0 | yes |  | unsolved |  |  |  |  |  |  |  |  |
| SI04 | non-PCD | M/23 | -1 | L | 1 | no |  | unsolved |  |  |  |  |  |  |  |  |
| SI05 | non-PCD | M/27 | 0.9 | R | 0 | no |  | unsolved |  |  |  |  |  |  |  |  |
| SI07 | non-PCD | F/35 | 0.9 | R | 1 | ? |  | unsolved |  |  |  |  |  |  |  |  |
| SI09 | non-PCD | F/36 | 0.7 | L<sup>§</sup> | 0 | no |  | unsolved |  |  |  |  |  |  |  |  |
| SI12 | non-PCD | F/40 | 0.9 | R | 0 | no | chet | *DNAH5* | Kartagener | None | 13900415 | A | C | - | E/* | stop gained |
|  |  |  |  |  |  |  | chet | *DNAH5* | Kartagener | None | 13769691 | G | GC | 0.00002 | A/X | frameshift |
| SI14 | non-PCD | M/18 | -0.8 | L | 1 | no |  | unsolved |  |  |  |  |  |  |  |  |
| SI16 | non-PCD | M/21 | -1 | L | 0 | no | hom | *CCDC151* | Kartagener | rs765121016 | 11533429 | T | TCTC | 0.00011 | E/- | inframe deletion |
| SI06 | PCD | M/46 | 1 | R | 0 | yes | hom | *LRRC6* | Kartagener | rs767624733 | 133687728 | T | C | 0.00017 | - | splice donor |
| SI08 | PCD | F/23 | 0.9 | R | 0 | yes | hom | *DNAH11* | PCD | rs373706559 | 21659620 | A | C | 0.00006 | Y/* | stop gained |
| SI11 | PCD | F/32 | 0.9 | R | 0 | yes | chet | *DNAAF1* | Kartagener | rs569633512 | 84203963 | C | T | - | - | splice donor |
|  |  |  |  |  |  |  | chet | *DNAAF1* | Kartagener | rs373103805 | 84193302 | G | C | 0.00023 | N/K | missense |
| SI13 | PCD | M/48 | 0.6 | L<sup>§</sup> | 0 | yes | hom | *CCDC114* | Kartagener | rs779459076 | 48814907 | CACG | C | 0.00093 | -/R | Inframe insert |
| SI15 | PCD | F/31 | 0.7 | R | 0 | yes | chet | *DNAH5* | Kartagener | None | 13786289 | A | T | - | K/* | stop gained |
|  |  |  |  |  |  |  | chet | *DNAH5* | Kartagener | rs397515540 | 13753397 | G | GA | 0.00034 | D/X | frameshift |
| SI17 | PCD | M/39 | 0.5 | R | 0 | yes | chet | *DNAH5* | Kartagener | rs548521732 | 13839638 | C | T | 0.00021 | - | splice acceptor |
|  |  |  |  |  |  |  | chet | *DNAH5* | Kartagener | rs769458738 | 13753597 | T | C | 0.00004 | R/H | missense |

Subjects shown with two mutations have possible compound heterozygous mutations, otherwise they have homozygous mutations. The Genome Reference Consortium (GRC) build 37 decoy version was used as reference sequence. EHI: Edinburgh Handedness Inventory score; NH: natural handedness; CHD: Congenital Heart Disease. Type: type of genetic mutation, i.e., homozygous (hom) or compound heterozygous (chet); MAF: minor allele frequency in population databases, if known; AAC: amino acid change. <sup>§</sup>Self-identified natural lefthander made to convert to right-handedness.

## Methods

### Participants

Fifteen people with radiologically documented SI, and 15 controls with normal *situs* matched for age, sex, education and handedness, were recruited from Ghent University Hospital and Middelheim Hospital Antwerp (approved by Research Ethics Committee). Table 1 gives an overview of the participants and their characteristics. All participants gave informed consent for DNA sample collection and genomic analysis in relation to body, brain and behavioural laterality. All methods were performed in accordance with the relevant guidelines and regulations.

Details about the recruitment, diagnosis and selection/exclusion procedure have been described previously ^18^. Radiological information (RX or CT) of thorax and complete abdomen was available in eight SI participants, and of thorax and upper abdomen in seven SI participants. The medical reports confirmed that all 15 participants presented with radiologically documented SI.

In five participants with SI, a formal diagnosis of PCD or Kartagener syndrome was found in their medical records. Kartagener syndrome refers to the clinical triad of situs inversus, chronic sinusitis, and bronchiectasis^10^. A sixth SI-participant was identified on account of a radiological consultation regarding infertility. The participant also presented with chronic sinusitis, mild chronic bronchitis. Although no formal diagnosis of PCD was obtained in this case, the presence of chronic upper and lower respiratory infection and infertility in an individual with SI warrants suspicion of Kartagener syndrome. Consequently, we included the participant within the PCD group. In addition, all six members of the PCD group reported having a daily wet cough, and had PICADAR scores of between 8 and 12 ^25^, and thus predictive probabilities of having PCD of between 66% and 99%, based on this recently-developed, questionnaire-based tool (note that five of these subjects anyway had formal medical diagnoses).

A seventh SI subject (SI03) had no medical record pertaining to PCD but did report daily wet cough. This subject had a PICADAR score of 8. In the study of Vingerhoets *et al.* this SI subject was classified as non-PCD, before the PICADAR assessment was available. For the purposes of the present study, we also treated this subject as non-PCD given the lack of formal medical history of PCD and the ambiguous PICADAR score, but we repeated some genetic analyses having excluded the person from the non-PCD group, in order to account for this uncertainty (see below).

Eight other SI subjects had no medical record of PCD or PCD-like symptoms, and were classified as non-PCD SI cases. Six of these reported no daily wet cough, and two did not answer in this regard. PICADAR scores can only be calculated in the presence of daily wet cough, so that none of these eight cases received PICADAR scores. Three of these cases had been previously diagnosed with congenital heart disease that required surgical treatment, and their radiological files all referred to their cardiac condition. Congenital heart disease is a frequent comorbidity of SI, as the cardiac circulation appears particularly sensitive to perturbation in normal left-right positional information ^26^.

Handedness was assessed using the Edinburgh Handedness Inventory (EHI) ^27^. Note that one non-PCD SI subject reported being forced to switch from left- to right-handedness in childhood, in which case five out of nine non-PCD SI cases were naturally left-handed. One of the six cases with PCD also reported forced left-to-right switching, otherwise the rest were right-handed (**Table 1**).

### Whole Genome Sequencing (WGS) and Pre-processing

DNA was extracted from saliva samples using the Oragene kit (Oragene). WGS was performed by Novogene (Hong Kong) using Illumina high throughput sequencing (HiSeq-PE150), creating paired end reads with a length of 150 base pairs (bp). Raw reads, stored in BAM files, were aligned to the human reference genome (the extended “decoy” version of b37) using Burrows-Wheeler Aligner (BWA) software ^28^, and sorted and reordered using SAMtools (v1.3.1) ^29^. PCR duplicates, which could arise during cluster amplification, were marked using Picard (v2.9.0). Genome Analysis Toolkit (GATK v3.7) ^30^ software was used to realign reads around insertions/deletions (indels) and to recalibrate base quality scores per sample.

### Variant Calling and Quality Control

Indels (insertion/deletions) and single nucleotide polymorphisms (SNPs) were called as recommended by the GATK Best Practices workflow. Variant Quality Score Recalibration (VQSR) was performed, and variants with a high probability of being false positive were flagged according to their sensitivity tranche (90, 99, 99.9 and 100). All SNPs and indels within a VQSR tranche of 99.9% or higher were discarded. Variants with a quality depth ≤ 9 or a call rate ≤ 0.8 were also removed. Vcftools v1.13 was used to create a summary table in the Variant Call Format (vcf). A total of 13,989,941 SNPs and indels were identified across the 30 subjects, with an average number of 5,186,055 (min= 5,053,188, max
 = 5,272,561) alternative alleles per subject (i.e. different to the reference genome, build37 decoy version).

### Genomic-level evaluation

For overall descriptive analysis of the participant genomes, a subset of 40,387 variants distributed genome-wide was used that had known Minor Allele Frequencies (MAFs) > 0.1, and were the result of pruning to be in low linkage disequilibrium with one another. For this, the flag --indep-pairwise function in Plink (v.1.90b3w) ^31^ was applied with a pairwise linkage disequilibrium r^2^ greater than 0.2, based on a SNP-SNP correlation matrix of 1500 by 150 in window size. The resulting data were then used as the basis for inferring pairwise identity by descent (IBD) sharing between subjects (i.e. genetic relatedness), inbreeding, and inconsistencies with reported sex, using the Plink operations --genome, --ibc/het, and --check-sex.

A different subset of 61,467 independent variants distributed genome-wide was used for visualizing population structure through Multi-Dimensional Scaling (MDS) analysis with respect to the 1000 genomes reference dataset (v37) ^32^ (downloaded from: ftp://climb.genomics.cn/pub/10.5524/100001_101000/100116/1kg_phase1_all.tar.gz) using the Plink operation –mds-plot.

### Annotation and filtering of single nucleotide variants affecting protein sequences

Variants were normalized using the software tool *vt normalize* (v0.5772-60f436c3) ^33^, which ensures consistent representation of variants across the genome. Normalized SNVs were annotated using Annovar ^34^ and Variant Effect Predictor (v88) ^35^. Gemini (v 0.20.0) was used to select protein coding variants with ‘MEDIUM’ or ‘HIGH’ impact severity annotations, as well as non-coding variants with ‘HIGH’ impact severity annotations (in practice those altering splice donor or acceptor sites). Additional filtering was done in R and comprised the removal of synonymous variants, and of ‘MEDIUM’ variants with a PolyPhen ^36^ or Sift ^37^ prediction score of “benign” or “tolerant” respectively. Data were then processed and analyzed separately under recessive and dominant models:

#### Recessive model

For the recessive model we further excluded variants with a known population frequency > 0.005 in any of the following databases: GNOMAD ^38^, ESP ^39^, 1000 Genomes ^32^ and ExAC databases ^38^. The remaining low-frequency variants were considered as putative mutations. Gene-level mutation counts per subject were then made, with a given gene being assigned as recessively mutated when it carried two copies of the same mutation (homozygous) or two different mutations (possible compound heterozygous). Integrative Genome Viewer (IGV v2.3.97) ^40^ was used to visualize the possible compound heterozygous mutations, and genes carrying these were discarded when both mutations were definitely present on the same allele (i.e. “in phase”) on a given sequence read. Finally, genes recessively mutated according to these criteria in any of the fifteen unaffected control subjects were excluded as being potentially causative in cases, and also removed for the purposes of Gene Set Enrichment Analysis (GSEA; see below): this step had the advantage of removing false variants from potential technical artifacts, or variants which are common in the population but not annotated as such in on-line databases. In total, 17 genes were excluded based on overlap with the unaffected control subjects. Genes on chromosome X were processed as part of the recessive pipeline, such that females would need to carry two mutations in a given gene, and males one mutation.

#### Dominant model

A maximum population frequency threshold of 5×10^−5^ was applied in this case, and genes carrying at least one rare variant according to this criterion were considered as potentially causative. Again, genes mutated according to this criterion in any of the fifteen unaffected control subjects (N = 47 genes) were excluded as being potentially causative in cases, and removed for the purposes of GSEA analysis (below).

### Gene Set Enrichment Analysis

To test whether a list of mutated genes in a given set of subjects contained functionally related genes across subjects, we performed Gene Set Enrichment Analysis (GSEA) using the g:Profiler R package (version 0.6.1) ^41^. A gene classification scheme derived from the Gene Ontology (GO) database ^42,43^ was used. This specified a total of 6380 functionally defined gene-sets, based on a background of 19,327 genes with functional annotations. Many genes are not represented in the GO, particularly mRNA transcripts of no known function or homology, so that the numbers of mutated genes in a given set of subjects was always higher than the subset used as input for GSEA.

Mutated genes on chromosome X were combined with recessively mutated autosomal genes for the purposes of GSEA. GSEAs were performed separately for the mutated gene lists in SI subjects with PCD (54 genes, of which 40 are in the GO), non-PCD SI subjects (60 genes, 38 in the GO), left handed non-PCD SI subjects (42 genes, 26 in the GO), and cases that were unsolved under the recessive model (42 genes, 22 in the GO). These analyses were also repeated after excluding subject SI03 (see above and **Table 1**), in the non-PCD SI group (55 genes, 35 in the GO), the left handed non-PCD SI group (37 genes, 23 in the GO), and in the unsolved group (36 genes, 19 in the GO).

For the dominant model, GSEA was performed separately for mutated genes in the non-PCD SI subjects (330 genes, 271 in the GO), the subset of non-PCD SI subjects that remained unsolved under the recessive model (217 genes, 175 in the GO), and the left handed subset of non-PCD subjects (201 genes, 163 in the GO). PCD subjects were not tested under a dominant model, as PCD is known to be a recessive phenotype. Dominant analyses were repeated after excluding subject SI03, for non-PCD SI subjects (285 genes, 235 in the GO), unsolved non-PCD SI subjects (170 genes, 139 in the GO) and non-PCD left handed SI subjects (156 genes, 127 in the GO).

In order to confirm that unaffected controls showed no significant pathway enrichment among their mutated genes, a mirrored exclusion was performed whereby any genes mutated in cases were excluded from the control gene lists. This resulted in 56 genes exclusively mutated in controls for the recessive model (of which 34 genes are in the GO), and 533 genes exclusively mutated in controls under the dominant model (440 in the GO).

The identities of genes were first converted using the g:Convert tool ^41^ to ensure recognition by the GO schema. The following settings were then used for GSEA: minimum set size = 15, maximum set size = 500, minimum intersect number = 2, hierarchical filtering = moderate. P-values were corrected for multiple testing across gene sets, based on the gSCS correction method in g:Profiler, separately for each input list of mutated genes corresponding to a given set of subjects. This method of multiple testing correction takes into account the hierarchical structure of the sets ^41^. We applied a cut-off P value of adjusted 0.01.

### Candidate Gene Lists

We queried the genetic data with respect to candidate gene lists for some purposes (see Results). An initial list of candidate genes was created in R (v3.3), based on searching the Online Mendelian Inheritance in Man (OMIM) database ^44^ for the terms: “situs inversus”, “heterotaxy | heterotaxia | situs ambiguus”, “congenital heart disease”, “PCD | ciliary dyskinesia | Kartagener”, “left-right”, and “asymmetry | laterality”. Additionally, the Clinvar column of our annotated variant data, which contains information based on the Clinvar database ^45^, was searched for these terms, and genes that were not yet in the initial list of candidate genes were accordingly included.

A broader candidate gene list was also created which included genes from the literature that were suggested to be associated with human laterality phenotypes and/or ciliopathies, but not (yet) present in our initial list of candidate genes (**Supplementary Table S1**). Specifically, a list of ciliopathy genes from a review of this topic ^6^ was searched for the terms: “PCD”, “heterotaxy”, “situs”, “left-right”, “asymmetry” or “laterality”, yielding 18 additional candidate genes. Additionally, a list of genes potentially associated with human laterality disorders, as compiled in a 2015 review ^2^, resulted in the addition of 25 candidate genes. Eleven additional genes were included, of which three were reported as potentially associated with PCD or heterotaxy ^46^, three had potential associations with non-syndromic heterotaxy ^47^, and five were considered as possibly causal in a recent exome sequencing study of various laterality defects ^48^. Finally, 39 more genes – arising from a search for the ‘situs inversus’ phenotype - were retrieved from the Mouse Genome Database (MGD) ^49^ at the Mouse Genome Informatics website, the Jackson laboratory, Bar Harbor, Maine. (URL: http://www.informatics.jax.org) [Oct, 2017].

### Structural Variants

For all participant genomes, structural Variants (SVs) (>50 kilobases in length) were called using a combination of two SV calling algorithms: CNVnator (v0.3.3) ^50^ and Lumpy (v 0.2.13) ^51^. These algorithms complement each other regarding the kinds of signals in WGS data that they can detect, as CNVnator is based on read-depth, whereas Lumpy is based on paired-end mapping ^52^. For CNVnator the bin size was set to 100 base pairs for all genomes except for two, where it was 150 base pairs (we determined a roughly optimal bin size for each subject’s genome, such that that the average read depth for that genome would be at least 4 standard deviations from zero). Lumpy was run using default parameters in “lumpyexpress”.

SVs were then annotated using SV2 ^53^. Variants that were present in any of the 15 healthy controls were removed from consideration as potentially causative for SI. Only variants that passed the default SV2 filtering criteria for quality were included ^53^.

## Results

### Protein-altering single nucleotide variants

#### Recessive mutations

Our variant calling, filtering and annotation pipeline produced between 5 and 15 recessively mutated genes per SI subject. We included six SI subjects with PCD as positive controls, in order to ensure that the variant calling and mutation definition criteria were well calibrated. As PCD is known to be a recessive phenotype for which at least 30 different genes have already been identified ^2^, we expected most, or all, of these six subjects to have identifiable mutations in known PCD-causing genes. As expected, each of these six cases had just one recessively mutated gene which was annotated ‘Kartagener’ or ‘PCD’ in the Clinvar database ^45^, and was therefore the most likely monogenic cause for their condition (**Table 1**). These genes were *LRRC6* [MIM:614930], *DNAH11* [MIM:603339], *DNAAF1* [MIM:613190], *CCDC114* [MIM:615038], and *DNAH5* [MIM: 603335] (the latter mutated in two SI cases with PCD, consistent with *DNAH5* being the most common cause of PCD in European-ancestry populations ^54^) (**Table 1**). The PCD subject with a homozygous *LRRC6* mutation (subject SI06) was the only individual to show an elevated inbreeding coefficient and non-European ancestry (**Supplementary Table S2, Supplementary Figure S1**).

None of the fifteen unaffected control subjects had any recessively mutated genes annotated ‘Kartagener’, ‘PCD’, ‘SA’ or ‘SI’ in Clinvar.

Surprisingly, two of the nine non-PCD SI cases had recessive mutations in genes annotated ‘Kartagener’ in the Clinvar database ^45^. These were subject SI12 (again involving mutations in *DNAH5*), and subject SI16 (in *CCDC151*) (**Table 1**). As these subjects had no medical records pertaining to PCD-like symptoms, and no daily wet cough, then the findings suggest reduced penetrance for PCD.

One of the nine non-PCD subjects – i.e., SI02 - had a recessive mutation in a gene that is annotated in Clinvar as ‘*situs ambiguus*’ and ‘*situs inversus totalis’*, but not annotated as PCD-causing (**Table 1**). This gene is *PKD1L1* [MIM: 617205]. A homozygous missense mutation in *PKD1L1* was previously reported in an individual with SI and congenital heart disease but no PCD, as well as recessive splicing mutations in two individuals with heterotaxy ^15^. Our subject SI02 had no diagnosis of congenital heart disease (CHD) (**Table 1**). As *PKD1L1* is a known recessive cause of non-PCD SI, we consider this gene to be the most likely cause of the non-PCD SI in subject SI02.

Subject SI03, who had no formal medical history of PCD, but had reported an intermediate PICADAR score, showed no recessive mutations in known PCD genes, which tends to support non-PCD status for this subject. This meant that six non-PCD SI cases did not have recessive mutations in genes known to cause human laterality disorders, as annotated in Clinvar, and therefore remained ‘unsolved’ under a recessive model (**Table 1**). Among these six non-PCD SI cases, four were left-handed, and were therefore of primary interest for the present study. Three of these same subjects also had CHD (**Table 1**).

We constructed an extended list of known or suspected laterality genes with reference to the literature and mouse laterality phenotypes (Methods; **Supplementary Table S1**), but none of these genes had recessive mutations in the six unsolved non-PCD SIT subjects.

We note possible compound heterozygous mutations in *PKD1* [MIM:601313], as a paralogue of *PKD1L1* [MIM:609721], in subject SI14 (**Supplementary Table S3**). However, mutations in this gene would be expected to cause autosomal dominant Polycystic Kidney Disease ^55,56^, and since there was no such diagnostic record for this subject, one or both of these specific mutations probably has limited functional impact and is therefore unlikely to be a monogenic cause for SI either.

*KIF13B* [MIM:607350] was putatively recessively mutated in subjects SI14 (unsolved) and SI12 (solved) (**Supplementary Table S3**). Although no literature has linked *KIF13B* to PCD or laterality phenotypes, *KIF3A* [MIM:604683], another kinesin encoding gene, is known to be involved in LR determination ^57^. Moreover, *KIF3B* [MIM: 603754] is known to affect motility of nodal cilia, accordingly affecting LR determination ^58^.Together with dyneins, kinesins allow ciliary proteins to enter the organelle via the transition zone by transporting them as cargo ^59,60^, and accordingly play an important role in ciliary construction and maintenance ^59,60^. The mutations in *KIF13B* might therefore potentially cause SI without PCD in subject SI14, and perhaps also affect the phenotypic outcome in subject SI12 who has likely causal mutations in *DNAH5*, but we cannot confidently assign a role to *KIF13B* on the basis of this evidence.

#### Dominant mutations

For the six non-PCD SI cases who remained unsolved under a recessive model, we also considered a dominant model using a maximum known mutation frequency of 5×10^−5^, and again cross-referenced the mutated genes against Clinvar and our extended candidate gene list (**Supplementary Table S1**), but no likely causative genes emerged (**Supplementary Table S3**) (see Discussion).

Subject SI05 showed a heterozygous mutation in *LRRC6* [MIM:614930] **(Supplementary Table 3)**, which was included among our candidate genes. However, since recessive – but not dominant - mutations in this gene have been associated with PCD ^61^, it is unlikely that this mutation is causal for non-PCD SI in this subject.

We also note a heterozygous mutation in *WDR62* [MIM:613583] in the unsolved case SI09 **(Supplementary Table 3)**. Although mice with mutations in this gene have shown dextrocardia and right pulmonary isomerism (MGI:5437081) ^49^, humans with recessive *WDR62* mutations do not show altered laterality. Instead, they have shown infantile spasm, microcephaly and intellectual disability ^62^. It is therefore unlikely that *WDR62* is a dominant cause of altered laterality in humans.

The gene *CEP290* [MIM:610142], mutated in subject SI14 **(Supplementary Table 3)**, is linked to left sided isomerism in mice (MGI:5438068)^49^. In humans, recessive mutations have been linked to a variety of ciliopathies, ranging from nephronophthisis, retinal degeneration and Joubert syndrome, to Bardet-Biedl syndrome and Meckel-Grüber syndrome ^63,64^. However, similar to the aforementioned genes, mutations in *CEP290* have not been associated with laterality phenotypes in humans, and we therefore consider this to be an unlikely cause of non-PCD SI.

Subject SI09 had a heterozygous mutation in PLXND1 (Supplementary Table 3), a gene which appears among search results for the phenotype ‘situs inversus’ within the Mouse Genome Database ^49^. However, while PLXND1 is annotated as a cause of aortic arch and atrial abnormalities in this database, it is not annotated with situs inversus or heterotaxia, so that the basis for the search result is unclear. We did not find evidence for this gene’s involvement in visceral laterality in a further literature search.

### Gene set enrichment analysis

We first performed gene set enrichment analysis using the positive control set of six SI subjects with PCD. As noted above, the six PCD subjects had likely recessive monogenic causes in five different genes (two subjects had mutations in the same gene, *DNAH5*). As expected, from the total combined list of recessively or X-linked mutated genes in these subjects, of which 40 genes were included in the GO schema, we obtained significant results for various cilia-related pathways, such as ‘axoneme’ (p = 0.004), ‘outer dynein arm assembly’ (p = 3.85×10^−5^), ‘dynein complex’ (p = 4.89×10^−5^), and ‘microtubule based movement’ (p = 1.41 ×10^−5^) (**Table 2**) (all P values adjusted for multiple testing across gene sets, see Methods). As expected, when the single most likely causative gene for each PCD subject was removed from the gene list, which left 36 recessively or X-linked mutated genes in these subjects that are present in the GO schema, there were no longer any significant enrichment terms, which further supports that the monogenic causes had been correctly identified in these subjects. The gene set enrichment analysis in the PCD subjects confirmed that, despite a relatively small number of subjects (i.e. six), the analysis was well powered to identify affected biological processes, even when most individual subjects had monogenic causes in different genes, and each subject had other, non-causative mutated genes.

**Table 2.** Gene set enrichment analysis under a recessive mutation model

| Group | p-value<sup>1</sup> | term size | query size<sup>2</sup> | overlap size | term ID | term name | intersection | samples | Pval Fisher<sup>3</sup> |
| --- | --- | --- | --- | --- | --- | --- | --- | --- | --- |
| Non-PCD SI (n=9)<sup>4</sup> | NS |  | 38 |  |  |  |  |  |  |
| Non-PCD SI LH (n=5)<sup>4</sup> | NS |  | 26 |  |  |  |  |  |  |
| Unsolved cases (n=6)<sup>4</sup> | NS |  | 22 |  |  |  |  |  |  |
| SI with PCD (n=6) | 3.85×10<sup>-5</sup> | 17 | 40 | 4 | GO:0036158 | outer dynein arm assembly | DNAH5, CCDC114, LRRC6, DNAAF1 | SI06, SI11, SI12, SI13, SI15, SI17 | 7.39×10<sup>-13</sup> |
| SI with PCD (n=6) | 1.41×10<sup>-4</sup> | 296 | 40 | 8 | GO:0007018 | microtubule-based movement | DNAH5, CCDC114, DNAH11, WDR60, LRRC6, DNAAF1, FMN2, DNAH12 | SI06, SI08, SI11, SI12, SI13, SI15, SI17 | 8.94×10<sup>-15</sup> |
| SI with PCD (n=6) | 4.89×10<sup>-5</sup> | 48 | 40 | 5 | GO:0030286 | dynein complex | DNAH5, CCDC114, DNAH11, WDR60, DNAH12 | SI08, SI11, SI12, SI13, SI15, SI17 | 1.59×10<sup>-13</sup> |
| SI with PCD (n=6) | 0.004 | 113 | 40 | 5 | GO:0005930 | axoneme | DNAH5, CCDC114, DNAH11, DNAAF1, RP1L1 | SI03, SI08, SI11, SI12, SI13, SI15, SI17 | 1.31×10<sup>-11</sup> |
| SI with PCD (n=6) | 0.002 | 427 | 40 | 8 | GO:0099568 | cytoplasmic region | DNAH5, CCDC114, DNAH11, SHROOM2, DNAAF1, FMN2, RP1L1, PCLO | SI03, SI06, SI08, SI11, SI12, SI13, SI15, SI17 | 1.73×10<sup>-13</sup> |
| Unaffected controls (n=15) | NS |  | 34 |  |  |  |  |  |  |

LH: left-handed.<sup>1</sup> P-values are corrected for multiple testing using the gSCS method. NS: no significant gene sets. <sup>2</sup> The number of mutated genes present in the GO schema.
<sup>3</sup>Uncorrected P-values were calculated using Fisher's exact test. <sup>4</sup> Results were the same after excluding subject SI03.

With this in mind, we then performed gene set enrichment analysis in the non-PCD SI cases, who were of primary interest for the present study due to a potential link with left-handedness. However, no significant enrichments were found when testing the list of recessively or X-linked mutated genes in the nine subjects with non-PCD SI, of which 38 genes were included in the GO schema. There was also no significant functional enrichment when testing the recessively mutated or X-linked genes in the five left-handed subjects with non-PCD SI, of which 26 genes were in the GO schema, and neither when testing the list of genes in the six unsolved non-PCD SI subjects, of which 22 genes were present in the GO schema (**Table 2**). Repeating the analysis after excluding subject SI03 from these subsets made no difference, there were still no significant gene sets. In addition, gene-set enrichment analysis with dominantly mutated genes as input did not produce significant results in the non-PCD SI cases, nor the left-handed or unsolved subsets.

As expected, the lists of recessively/X-linked and dominantly mutated genes in the fifteen unaffected control subjects did not produce any significant gene set enrichment terms (**Table 2**).

### Structural variant analysis

We additionally screened the genomes of all subjects for structural variants (such as larger-scale deletions and duplications), using two complementary algorithms (see Methods). SI subjects had SVs affecting an average of 9.7 genes per subject (min = 2 SVs, max = 16 SVs), and controls had SVs affecting an average of 9.6 genes per subject (min = 5 SVs, max = 16 SVs). None of the SI subjects, regardless of PCD status, had SVs affecting genes that were annotated SI, PCD, Kartagener, *situs ambiguus* (SA), or Heterotaxy (HTX) in Clinvar, nor affecting genes from our broader list of candidate laterality genes.

## Discussion

In this study we aimed to identify rare, highly penetrant genetic mutations that might link visceral body asymmetry with handedness, by analysing whole genome sequence data from nine SI subjects without medical histories of PCD, five of whom were left-handed. We additionally included six SI subjects with PCD as positive technical controls, and fifteen unaffected subjects as negative controls.

Likely monogenic causes were identified in all positive controls, i.e. each of the six PCD subjects had one recessively mutated gene (usually a different gene in each subject) that is already known to cause PCD when mutated. The six PCD subjects also confirmed that significant pathway enrichment could be detected on the basis of their mutated gene lists, as multiple gene sets related to ciliary functions were detected. The fifteen unaffected control subjects showed no recessive mutations in genes known to cause PCD or laterality phenotypes, as expected.

Two of the nine non-PCD SI subjects had likely recessive monogenic causes in known PCD genes. This may indicate reduced penetrance of these mutations for PCD-like symptoms such as lung symptoms, although they can apparently affect *situs* in early development. One non-PCD SI subject, who was right-handed, had a homozygous mutation in a gene already known to cause SI without PCD, i.e. *PKD1L1* ^15^. This gene encodes a component of a calcium channel which is associated with non-motile cilia ^65^.

However, the six remaining non-PCD SI subjects had no obvious monogenic basis for their condition, i.e. they did not have likely causative mutations in genes known to cause human laterality disorders as annotated in the Clinvar database, nor within an extended list of known or suspected laterality genes based on literature searching and mouse phenotypes. Among the six non-PCD SI subjects, four were left-handed, and therefore comprised the bulk of left-handers in the dataset. Furthermore, gene set enrichment analysis of their mutated genes did not identify significant biological pathways, in either the whole set, or the left-handed subset, or the subset that was ‘unsolved’ by recessive monogenic causes. In other words, the biology of their non-PCD SI could not be linked via the mutations that they carried. Finally, we also considered larger genomic rearrangements known as Copy Number Variants (CNVs), but no obvious candidate genes emerged.

A monogenic model is still possible for the majority of the non-PCD SI cases, and/or for the left-handed subset specifically, but would have to involve genetic heterogeneity across a set of genes which are not currently linked in terms of their known biology, at least to an extent which would have been discernible in this dataset, as it was for the PCD subjects. We did not therefore identify a genetic-developmental pathway that links handedness and visceral asymmetry in this study. The question of whether, and how, functional brain laterality is linked developmentally to visceral laterality in humans remains unanswered ^20^.

Genetic contributions to non-PCD SI and left-handedness might also involve non-coding variation, or rare combinations of multiple common variants. The noncoding genome comprises 98% of the genome, but interpreting the variation within these regions is challenging. Several attempts have been made to rank potentially causative variants across the genome based on scores that integrate different types of information, including conservation of DNA sequence, regulatory information ^66^, and population genomic data ^67-72^. However, these ranking approaches are currently not very concordant with each other ^73^. For the present study we did not address these possibilities, which must await larger sample sizes and an improved understanding of the role of rare, non-coding mutations in phenotypic variation.

*In utero* environmental effects such as prenatal drug exposure might also affect left-right determination of body and brain ^74^. Handedness itself is known to associate with various early life factors including birthweight and breastfeeding, although not to a degree which is remotely predictive at the individual level ^75^. As noted in the introduction, left-handedness has a modest heritability of roughly 25% in family and twin studies, and lower in SNP-based heritability studies. Regardless, the primary causes of this trait remain unknown.

In this study there was a degree of diagnostic uncertainty as regards the PCD status of some SI subjects. However, it was not the purpose of the present study to achieve a clinical diagnosis of the presence or absence of PCD, nor to confirm already-known PCD genes. In this context we did not, for example, sequence the mutations in the PCD subjects by another technique for validation, nor confirm allelic phase in the compound heterozygotes. Rather, we were concerned to identify potentially causative mutations in the nine SI subjects without medical histories of PCD who show an elevated rate of left-handedness, with the potential to yield important basic insights into body and brain laterality. If we had found clear candidate mutations in left-handed members of the non-PCD SI group, in genes not previously linked to PCD, then further validation and functional characterisation of those mutations would have been appropriate, but this did not arise.

Regardless of the PCD status of any individual SI subject in this study, the overall pattern of results was clearly different between the PCD and non-PCD groups, where all six positive control subjects in the PCD group had mutations in known PCD-causing genes, while six out of nine in the non-PCD group had no obvious monogenic mutations, among whom were most of the left-handers. Also, the pathway analyses in various different subsets of the non-PCD group showed a consistent lack of significant enrichment, whereas clear signals emerged from the PCD group, which further supports an overall distinction of the groups. Nonetheless, further detailed investigation of subjects SI03, SI12 and SI16 with PCD diagnostic tools might reveal lung and other ciliary symptoms to an extent ^76^.

Although ciliary-induced nodal flow plays a crucial role in symmetry breaking, some organisms, such as chicks and fruit flies (*Drosophilia melanogaster*), do not have motile nodal cilia, yet they do show asymmetrical organs ^77^. For example, left-right asymmetry in chicks involves rearranging the relative orientations of cells expressing critical genes at the node, in a non-ciliary-dependent manner ^78^. Furthermore, left-right asymmetry of the *Drosophilia* gut is established by intracellular cytoskeletal organization that may give rise to cellular shape chirality, by means of unidirectional tilting of radial fibers, and anti-clockwise swirling of transverse fibers ^79^. Whether such mechanisms also influence left-right organ asymmetry in mammals is unclear. In the present study we saw no mutations in the homologues of two genes implicated in cellular chirality in *Drosophilia, MYO1D* or *MYO1C* ^80^, nor in the homologues of two genes thought to affect laterality in chicks and frogs, *SLC6A4* and *SLC18A2* ^81^

Future studies in larger human cohorts may help to identify genetic contributions to non-PCD SI and left-handedness in some cases. Candidate biological pathways which emerge from research on non-mammalian mechanisms of asymmetry development should be considered in future studies.

### Data availability

Requests to access the genomic datasets generated for the current study will be considered in relation to the consents, relevant rules and regulations, and can be made via the corresponding author.

## Acknowledgements

Thanks to all of the study participants. This research was funded by the Max Planck Society (Germany) and the Netherlands Organization for Scientific Research (NWO, grant No. 054-15-101) as part of the FLAG-ERA consortium project ‘MULTI-LATERAL’, a Partner Project to the European Union’s Flagship Human Brain Project. The study was also funded by the Fonds Wetenschappelijk Onderzoek-Vlaanderen (FWO-grant n^0^ G.0114.16N) assigned to GV.

## Author contributions

GV recruited and characterized the subjects. GV, SEF & CF conceived the study. MCP performed the genetic data analysis with support from ACC. MCP and CF wrote the manuscript. GV, ACC & SEF edited the manuscript. CF directed the study.

## Additional information

The authors declare no competing interests.

## Notes

**Conflicts of interest** The authors declare no conflicts of interest.

