## Supplementary for "No clear monogenic links between left-handedness and *situs inversus*"

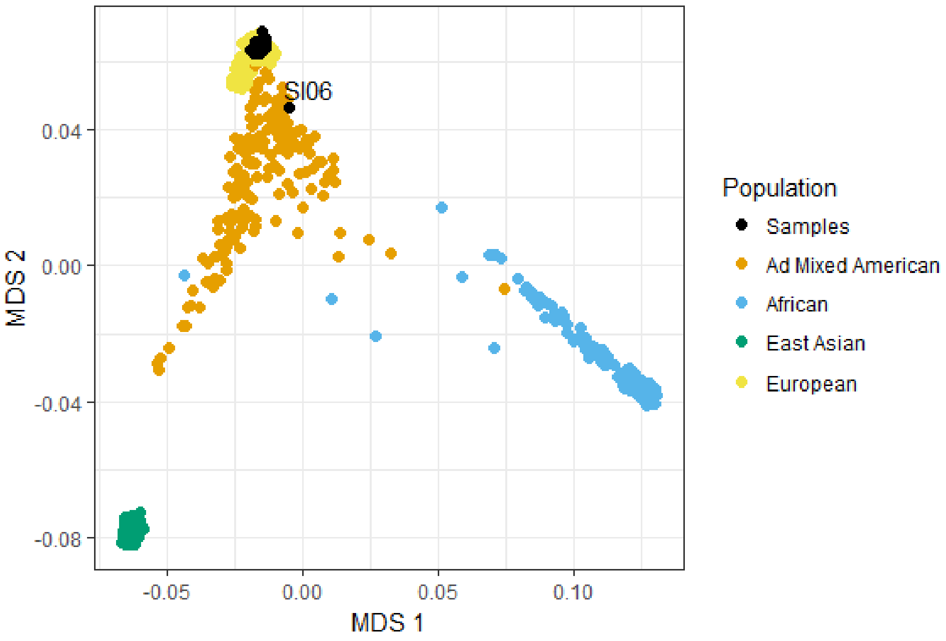

**Figure S1**. Multidimensional Scaling (MDS) to capture overall genomic diversity among the 30 study samples (black dots), in relation to the 1000 Genomes populations of known geographic ancestries.Subject SI06 is clearly distinct from the European-descent population (yellow dots).

**Table S1.** Broader list of candidate genes, and the sources that led to their inclusion.

| Gene | Location | Ensembl Gene ID | Literature |
|---|---|---|---|
| AK7 | 14q32.2 | ENSG00000140057 | Reiter & Leroux (2017) |
| AMN | 14q32 | ENSG00000166126 | Mouse Genome Database (2018) |
| ANKS3 | 16p13.3 | ENSG00000168096 | Reiter & Leroux (2017) |
| ANKS6 | 9q22.33 | ENSG00000165138 | Reiter & Leroux (2017) |
| AP1B1 | 22q12 | ENSG00000100280 | Mouse Genome Database (2018) |
| ATMIN | 16q23.2 | ENSG00000166454 | Mouse Genome Database (2018) |
| BBS1 | 11q13 | ENSG00000174483 | Mouse Genome Database (2018) |
| BBS2 | 16q21 | ENSG00000125124 | Deng, Xia, & Deng (2015) |
| BBS4 | 15q22.3-q23 | ENSG00000140463 | Mouse Genome Database (2018) |
| BICC1 | 10q21.2 | ENSG00000122870 | Mouse Genome Database (2018) |
| BMS1 | 10q11.21 | ENSG00000165733 | Li et al. (2019) |
| CC2D2A | 4p15.3 | ENSG00000048342 | Mouse Genome Database (2018) |
| CEP290 | 12q21.3 | ENSG00000198707 | Mouse Genome Database (2018) |
| CFAP54 | NA | NA | Reiter & Leroux (2017) |
| CITED2 | 6q23.3 | ENSG00000164442 | Deng, Xia, & Deng (2015) |
| DAND5 | 19p13.2-p13.13 | ENSG00000179284 | Mouse Genome Database (2018) |
| DCTN5 | 16p12.2 | ENSG00000166847 | Mouse Genome Database (2018) |
| DNAH6 | 2p11.2 | ENSG00000115423 | Reiter & Leroux (2017) |
| DNAH9 | 17p12 | ENSG00000007174 | Reiter & Leroux (2017) |
| DPCD | 10q24.32 | ENSG00000166171 | Reiter & Leroux (2017) |
| DRC3 | NA | NA | Reiter & Leroux (2017) |
| EPB41L5 | 2q14 | ENSG00000115109 | Deng, Xia, & Deng (2015) |
| FGF10 | 5p13-p12 | ENSG00000070193 | Mouse Genome Database (2018) |
| FOXA2 | 20p11 | ENSG00000125798 | Peeters & Devriendt (2006) |
| FOXH1 | 8q24.3 | ENSG00000160973 | Deng, Xia, & Deng (2015) |
| FOXI2 | 10q26.2 | ENSG00000186766 | Hagen et al. (2016) |
| FOXJ1 | 17q22-q25 | ENSG00000129654 | Mouse Genome Database (2018) |
| FOXL2 | 3q23 | ENSG00000183770 | Mouse Genome Database (2018) |
| GALNT11 | 7q36.1 | ENSG00000178234 | Reiter & Leroux (2017) |
| GDF1 | 19p12 | ENSG00000223802 | Deng, Xia, & Deng (2015) |
| GJA1 | 5q22.31 | ENSG00000152661 | Hagen et al. (2016) |
| HES7 | 17p13.2 | ENSG00000179111 | Deng, Xia, & Deng (2015) |
| IFT122 | 3q21 | ENSG00000163913 | Mouse Genome Database (2018) |
| IFT140 | 16p13.3 | ENSG00000187535 | Mouse Genome Database (2018) |
| IFT27 | 22q12.3 | ENSG00000100360 | Mouse Genome Database (2018) |
| IFT74 | 9p21.2 | ENSG00000096872 | Mouse Genome Database (2018) |
| IFT88 | 13q12.1 | ENSG00000032742 | Reiter & Leroux (2017) |
| IHH | 2q33-q35 | ENSG00000163501 | Mouse Genome Database (2018) |
| INVS | 9q31 | ENSG00000119509 | Reiter & Leroux (2017) |
| ISL1 | 5q | ENSG00000016082 | Li et al. (2019) |
| KIF3A | 5q31 | ENSG00000131437 | Reiter & Leroux (2017) |
| LEFTY1 | 1q42.1 | ENSG00000243709 | Mouse Genome Database (2018) |
| LETYA | NA | NA | Peeters & Devriendt (2006) |
| LZTFL1 | 3p21.3 | ENSG00000163818 | Deng, Xia, & Deng (2015) |
| MBD4 | 3q21-q22 | ENSG00000129071 | Mouse Genome Database (2018) |
| MCIDAS | 5q11.2 | ENSG00000234602 | Reiter & Leroux (2017) |
| MED13L | 12q24 | ENSG00000123066 | Deng, Xia, & Deng (2015) |

| Gene | Location | Ensembl Gene ID | Literature |
|---|---|---|---|
| MEGF8 | 19q12 | ENSG00000105429 | Deng, Xia, & Deng (2015) |
| MGAT1 | 5q35 | ENSG00000131446 | Mouse Genome Database (2018) |
| MGRN1 | 16p13.3 | ENSG00000102858 | Mouse Genome Database (2018) |
| MKS1 | 17q23 | ENSG00000011143 | Mouse Genome Database (2018) |
| MNS1 | 15q21.3 | ENSG00000138587 | Reiter & Leroux (2017) |
| NEK2 | 1q32.2-q41 | ENSG00000117650 | Deng, Xia, & Deng (2015) |
| NEK8 | 17q11.1 | ENSG00000160602 | Deng, Xia, & Deng (2015) |
| NKX2-5 | 5q34 | ENSG00000183072 | Deng, Xia, & Deng (2015) |
| NKX2.5 | NA | NA | Peeters & Devriendt (2006) |
| NKX6-2 | 10q26 | ENSG00000148826 | Hagen et al. (2016) |
| NME7 | 1q24 | ENSG00000143156 | Mouse Genome Database (2018) |
| NOTCH1 | 9q34.3 | ENSG00000148400 | Deng, Xia, & Deng (2015) |
| NOTCH2 | 1p13-p11 | ENSG00000134250 | Deng, Xia, & Deng (2015) |
| NOTO | NA | NA | Reiter & Leroux (2017) |
| NPHP3 | 3q22 | ENSG00000113971 | Deng, Xia, & Deng (2015) |
| NPHP4 | 1p36 | ENSG00000131697 | Deng, Xia, & Deng (2015) |
| NUP188 | 9q34.11 | ENSG00000095319 | Deng, Xia, & Deng (2015) |
| OFD1 | Xp22.3-p22.2 | ENSG00000046651 | Reiter & Leroux (2017) |
| PAX8 | 2q12-q14 | ENSG00000125618 | Mouse Genome Database (2018) |
| PCSK5 | 9q21.3 | ENSG00000099139 | Mouse Genome Database (2018) |
| PCSK6 | 15q26 | ENSG00000140479 | Mouse Genome Database (2018) |
| PITX2 | 4q25-q26 | ENSG00000164093 | Deng, Xia, & Deng (2015) |
| PKD2 | 4q21-q23 | ENSG00000118762 | Deng, Xia, & Deng (2015) |
| PLXND1 | 3q22 | ENSG00000004399 | Mouse Genome Database (2018) |
| POLB | 8p11.2 | ENSG00000070501 | Mouse Genome Database (2018) |
| PSKH1 | 16q22.1 | ENSG00000159792 | Mouse Genome Database (2018) |
| PXDNL | 8q11 | ENSG00000147485 | Li et al. (2019) |
| RAI2 | Xp22 | ENSG00000131831 | Li et al. (2019) |
| RFX3 | 9p24.2 | ENSG00000080298 | Reiter & Leroux (2017) |
| RIPPLY1 | Xq22.3 | ENSG00000147223 | Li et al. (2019) |
| ROCK2 | 2p24 | ENSG00000134318 | Deng, Xia, & Deng (2015) |
| RPGR | Xp11.4 | ENSG00000156313 | Reiter & Leroux (2017) |
| RPGRIP1L | 16q12.2 | ENSG00000103494 | Mouse Genome Database (2018) |
| SESN1 | 6q21 | ENSG00000080546 | Deng, Xia, & Deng (2015) |
| SHH | 7q36 | ENSG00000164690 | Mouse Genome Database (2018) |
| SHROOM3 | 4q21.1 | ENSG00000138771 | Deng, Xia, & Deng (2015) |
| SLIT2 | 4p15.2 | ENSG00000145147 | Mouse Genome Database (2018) |
| SMAD2 | 18q21 | ENSG00000175387 | Deng, Xia, & Deng (2015) |
| T | 6q27 | ENSG00000164458 | Mouse Genome Database (2018) |
| TBC1D32 | 6q22.31 | ENSG00000146350 | Mouse Genome Database (2018) |
| TGFBR2 | 3p22 | ENSG00000163513 | Deng, Xia, & Deng (2015) |
| TGIF1 | 18p11.3 | ENSG00000177426 | Mouse Genome Database (2018) |
| TMEM67 | 8q21.13-q22.1 | ENSG00000164953 | Mouse Genome Database (2018) |
| TRAPPC10 | 21q22.3 | ENSG00000160218 | Mouse Genome Database (2018) |
| UVRAG | 11q13 | ENSG00000198382 | Deng, Xia, & Deng (2015) |
| WDR62 | 19q13.12 | ENSG00000075702 | Mouse Genome Database (2018) |

**Table S2.** Inbreeding coefficients per subject.

| Subject | Group | O(HOM) | E(HOM) | NOMISS | Fhat1 | Fhat2 | Fhat3 |
|---|---|---|---|---|---|---|---|
| SI02 | non-PCD SI | 21098 | 21680 | 40351 | -0.034 | -0.034 | -0.034 |
| SI03 | non-PCD SI | 20997 | 21680 | 40352 | -0.035 | -0.038 | -0.036 |
| SI04 | non-PCD SI | 21099 | 21680 | 40350 | -0.027 | -0.035 | -0.031 |
| SI05 | non-PCD SI | 21161 | 21680 | 40337 | -0.026 | -0.030 | -0.028 |
| SI07 | non-PCD SI | 20935 | 21690 | 40354 | -0.047 | -0.039 | -0.043 |
| SI09 | non-PCD SI | 21090 | 21690 | 40363 | -0.037 | -0.033 | -0.035 |
| SI12 | non-PCD SI | 21274 | 21690 | 40359 | -0.032 | -0.023 | -0.028 |
| SI14 | non-PCD SI | 20952 | 21690 | 40359 | -0.041 | -0.040 | -0.040 |
| SI16 | non-PCD SI | 21282 | 21680 | 40352 | -0.018 | -0.023 | -0.021 |
| **SI06** | **SI with PCD** | **22503** | **21680** | **40348** | **0.048** | **0.041** | **0.044** |
| SI08 | SI with PCD | 21039 | 21690 | 40357 | -0.033 | -0.035 | -0.034 |
| SI11 | SI with PCD | 21059 | 21680 | 40346 | -0.036 | -0.034 | -0.035 |
| SI13 | SI with PCD | 21011 | 21690 | 40360 | -0.036 | -0.036 | -0.036 |
| SI15 | SI with PCD | 21000 | 21690 | 40355 | -0.032 | -0.038 | -0.035 |
| SI17 | SI with PCD | 20888 | 21690 | 40363 | -0.042 | -0.047 | -0.045 |
| CO02 | Unaffected control | 21221 | 21690 | 40358 | -0.031 | -0.025 | -0.028 |
| CO03bis | Unaffected control | 21028 | 21680 | 40341 | -0.036 | -0.036 | -0.036 |
| CO04 | Unaffected control | 21192 | 21680 | 40351 | -0.026 | -0.030 | -0.028 |
| CO15 | Unaffected control | 21177 | 21680 | 40348 | -0.037 | -0.025 | -0.031 |
| CO06bis | Unaffected control | 21151 | 21680 | 40346 | -0.035 | -0.029 | -0.032 |
| CO07 | Unaffected control | 20989 | 21680 | 40349 | -0.038 | -0.038 | -0.038 |
| CO08 | Unaffected control | 21128 | 21680 | 40354 | -0.034 | -0.030 | -0.032 |
| CO09bis | Unaffected control | 20923 | 21680 | 40350 | -0.039 | -0.045 | -0.042 |
| CO11 | Unaffected control | 21095 | 21680 | 40344 | -0.038 | -0.029 | -0.033 |
| CO12 | Unaffected control | 21095 | 21690 | 40356 | -0.031 | -0.034 | -0.032 |
| CO13bis | Unaffected control | 21067 | 21690 | 40358 | -0.039 | -0.033 | -0.036 |
| CO14 | Unaffected control | 20983 | 21690 | 40362 | -0.041 | -0.039 | -0.040 |
| CO15 | Unaffected control | 20965 | 21690 | 40360 | -0.036 | -0.042 | -0.039 |
| CO16bis | Unaffected control | 21007 | 21690 | 40356 | -0.038 | -0.038 | -0.038 |
| CO17 | Unaffected control | 21170 | 21680 | 40345 | -0.023 | -0.031 | -0.027 |

O(HOM): observed number of homozygotes; E(HOM): expected number of homozygotes; NOMISS: Number of non-missing genotype calls; Fhat1: variance-standardized relationship minus 1; Fhat2: Excess homozygosity-based inbreeding estimate; Fhat3: estimate based on correlation between uniting gametes. Subject SI06 (bold font) is the only subject with positive inbreeding coefficients, indicating an excess of homozygosity

**Table S3**. Notable mutations in unsolved cases, which we nonetheless do not consider causative for their SI (see the Results section of the main text for explanation).

| Subj | SI group | Sex/ Age | EHI | NH | CHD | Daily wet cough | Type | Gene | Clinvar annotation for the gene | rs ID | Start position | ref | alt | MAF | AAC | impact |
|---|---|---|---|---|---|---|---|---|---|---|---|---|---|---|---|---|
| SI03 | non-PCD | F/26 | -0.8 | L | 0 | yes | dom | PKD1 | 3-4 toe syndactyly \| abnormality of the kidney \| cerebral aneurysm \| hepatic cysts \| hereditary cancer-predisposing syndrome \| hypertension \| inborn genetic diseases \| lymphangiomyomatosis \| moderate sensorineural hearing impairment \| multicystic kidney dysplasia \| multiple renal cysts \| pancreatic cysts \| polycystic kidney disease\x2c adult type \| polycystic kidney dysplasia \| proteinuria \| renal cyst \| renovascular hypertension \| stage 5 chronic kidney disease \| tuberous sclerosis 2 \| tuberous sclerosis and lymphangiomyomatosis \| tuberous sclerosis syndrome | None | 2163259 | T | C | -1 | M/V | missense variant |
| SI05 | non-PCD | M/27 | 0.9 | R | 0 | no | dom | LRRC6 | Kartagener | None | 133687517 | A | T | -1 | NA | splice donor variant |
| SI09 | non-PCD | F/36 | 0.7 | L§ | 0 | no | dom | WDR62 | abnormality of neuronal migration \| | None | 36594252 | CT | C | -1 | L/X | frameshift variant |

| | | | | | | | | | | | | | | | | |
|---|---|---|---|---|---|---|---|---|---|---|---|---|---|---|---|---|
| | | | | | | | | | microcephaly\x2c cortical malformations\x2c and intellectual disability \| primary microcephaly\x2c recessive \| primary microcephaly 2 with or without cortical malformations \| primary autosomal recessive microcephaly 2 | | | | | | | |
| SI09 | non-PCD | F/36 | 0.7 | L§ | 0 | no | dom | PLXND1 | None | None | 129286636 | AGAC | A | 9.01E-06 | V/- | inframe deletion |
| SI12, SI14 | non-PCD | M/18 | -0.8 | L | 1 | no | chet | KIF13B | None | None | 28974427 | C | A | -1 | V/L | missense variant |
| SI12, SI14 | non-PCD | M/18 | -0.8 | L | 1 | no | chet | KIF13B | None | rs753108980 | 28974427 | C | T | 0.000188 | V/M | missense variant |
| SI14 | non-PCD | M/18 | -0.8 | L | 1 | no | chet | PKD1 | See above | rs199700485 | 2154530 | G | T | 0.001591 | T/N | missense variant |
| SI14 | non-PCD | M/18 | -0.8 | L | 1 | no | chet | PKD1 | See above | rs142733588 | 2153266 | C | T | 0.000245 | G/S | missense variant |
| SI14 | non-PCD | M/18 | -0.8 | L | 1 | no | dom | CEP290 | abnormality of the kidney \| agenesis of cerebellar vermis \| bardet-biedl syndrome \| bardet-biedl syndrome 14\|blindness\|cep290-related disorders \| central hypotonia \| cerebellar cyst \| cerebellar vermis hypoplasia \| familial aplasia of the vermis \| global developmental delay \| hyperechogenic kidneys \| joubert syndrome and related disorders \| joubert | None | 88462420 | C | A | -1 | A/S | missense variant |

| | | | | | | | | | |
|---|---|---|---|---|---|---|---|---|---|
| | | | | | | | | | syndrome 5\|leber congenital amaurosis \| leber congenital amaurosis 10\|meckel-gruber syndrome \| meckel syndrome type 4\|molar tooth sign on mri\|nephronophthisis \| nephronophthisis,not specified \| nystagmus \| polycystic kidney dysplasia \| renal dysplasia and retinal aplasia \| retinal dystrophy \| senior-loken syndrome 6 |

The Genome Reference Consorium (GRC) build 37 decoy version was used as reference sequence. EHI: Edinburgh Handedness Inventory score; NH: natural handedness; CHD: Congenital Heart Disease. Type: type of genetic mutation, i.e., heterozygous (dom), homozygous (hom) or compound heterozygous (chet); MAF: minor allele frequency in population databases, if known; AAC: amino acid change. [§]Self-identified natural lefthander made to convert to right-handedness.
